# Deducing high-accuracy protein contact-maps from a triplet of coevolutionary matrices through deep residual convolutional networks

**DOI:** 10.1101/2020.10.05.326140

**Authors:** Yang Li, Chengxin Zhang, Eric W. Bell, Wei Zheng, Xiaogen Zhou, Dong-Jun Yu, Yang Zhang

**Affiliations:** School of Computer Science and Engineering, Nanjing University of Science and Technology, Xiaolingwei 200, Nanjing, China, 210094; Department of Computational Medicine and Bioinformatics, University of Michigan, Ann Arbor, MI 48109 USA

## Abstract

The topology of protein folds can be specified by the inter-residue contact-maps and accurate contact-map prediction can help *ab initio* structure folding. We developed TripletRes to deduce protein contact-maps from discretized distance profiles by end-to-end training of deep residual neural-networks. Compared to previous approaches, the major advantage of TripletRes is in its ability to learn and directly fuse a triplet of coevolutionary matrices extracted from the whole-genome and metagenome databases and therefore minimize the information loss during the course of contact model training. TripletRes was tested on a large set of 245 non-homologous proteins from CASP and CAMEO experiments, and outperformed other state-of-the-art methods by at least 58.4% for the CASP 11&12 and 44.4% for the CAMEO targets in the top-*L* long-range contact precision. On the 31 FM targets from the latest CASP13 challenge, TripletRes achieved the highest precision (71.6%) for the top-*L*/5 long-range contact predictions. These results demonstrate a novel efficient approach to extend the power of deep convolutional networks for high-accuracy medium- and long-range protein contact-map predictions starting from primary sequences, which are critical for constructing 3D structure of proteins that lack homologous templates in the PDB library.

**Availability:** The training and testing data, standalone package, and the online server for TripletRes are available at https://zhanglab.ccmb.med.umich.edu/TripletRes/.

**Author Summary:** *Ab initio* protein folding has been a major unsolved problem in computational biology for more than half a century. Recent community-wide Critical Assessment of Structure Prediction (CASP) experiments have witnessed exciting progress on *ab initio* structure prediction, which was mainly powered by the boosting of contact-map prediction as the latter can be used as constraints to guide *ab initio* folding simulations. In this work, we proposed a new open-source deep-learning architecture, TripletRes, built on the residual convolutional neural networks for high-accuracy contact prediction. The large-scale benchmark and blind test results demonstrate significant advancement of the proposed methods over other approaches in predicting medium- and long-range contact-maps that are critical for guiding protein folding simulations. Detailed data analyses showed that the major advantage of TripletRes lies in the unique protocol to fuse multiple evolutionary feature matrices which are directly extracted from whole-genome and metagenome databases and therefore minimize the information loss during the contact model training.

## Introduction

Protein structure prediction represents an important unsolved problem in computational biology, with the major challenge on distant-homology modeling (or *ab initio* structure prediction)(1, 2). Recent CASP experiments have witnessed encouraging progress in protein contact predictions, which have been proven to be helpful to improve accuracy and success rate for distant-homologous protein targets(3-6).

The idea of developing sequence-based contact-map prediction to assist *ab initio* protein structure prediction is, however, not new, which can be traced back to at least 25 years ago(7, 8). In general, the methods for sequence-based protein contact-map prediction can be classified into two categories: coevolution analysis methods (CAMs) and machine learning methods (MLMs). In CAM, the predictors try to predict inter-residue contacts by analyzing evolutionary correlations of the target residue pairs from multiple sequence alignments (MSAs), under the assumption that correlated mutations in evolution usually correspond to spatial contacts of residue pairs. The CAMs can be further divided into local and global approaches. The local approaches use correlation coefficient, e.g., mutation information(7) and covariance(9), to predict contacts; these approaches are “local” because they predict contact between two residue positions regardless of other positions. In contrast, the global approaches, also called direct coupling analysis (DCA) methods, consider effects from other positions to better quantifying the strength of direct relationship between two residue positions. DCA models demonstrated significant advantage over the local approaches, and essentially re-stimulated the interest of the field of protein structure prediction in contact-map predictions. However, the success of most DCA methods(10-15) is still limited for the proteins with few sequence homologs, because a shallow MSA significantly reduce the accuracy of DCA to derive the inherent correlated mutations. In addition, DCA models only capture linear relationships between residues while residueresidue relationships in proteins are inherently non-linear.

As a more general approach, MLMs intend to learn the inter-residue contacts from sequential information and coevolution analysis features with supervised machine learning models trained with known structures from the PDB. Early attempts utilized support vector machines (SVMs)(16, 17), random forests (RFs)(12, 18), artificial neural networks (NNs)(19-22) etc., to model the complex relationships between residues. Recently, great improvements have been achieved by the application of convolutional neural networks (CNNs) in several predictors, including DNCON2(23), DeepContact(24) and RaptorX-Contact(25). Most of the predictors were however trained on the final contact-map confidence scores(23-25), which may suffer coevolutionary information loss in data postprocessing. In a recent study, we proposed ResPRE(26) which directly utilized the ridge-regularized precision matrices calculated from raw alignments without post-processing in regular coevolution analysis features. Although it uses the evolutionary matrix as the only input feature, the performance of ResPRE was comparable to many state-of-the-art methods that combine additional one-dimensional features, such as solvent accessibility, predicted secondary structure and physicochemical properties. Despite the success, ResPRE still bears several shortcomings. First, ResPRE lacks consideration for multiple coevolutionary matrices as features, which could provide complementary information. Second, it was trained by the supervision of binary protein contact-maps that lack continuous inter-residue distance information. Finally, the coevolution features were derived from a somewhat simplified HHblits(27) MSA collection procedure, which did not always include sufficient homologous sequences for meaningful precision matrix generation.

In this work, we proposed a new deep learning architecture, TripletRes, built on a residual neural network protocol(28) to integrate a triplet of coevolutionary matrices features from pseudolikelihood maximization of Potts model, precision matrix and covariance matrix for high-accuracy contact-map prediction (Fig 1). The model was trained on a non-redundant subset of sequences with known PDB structures supervised by discretized inter-residue distance-maps in order to capture the inherent distance information between residues, where a new deep MSA generation protocol was employed to derive the coevolutionary matrices. The benchmark results on the public CASP and CAMEO targets, along with the community-wide blind tests in the most recent CASP13 experiment, show that the new approach is capable to create contact-maps with precision significantly beyond previous methods.

**Fig 1.**
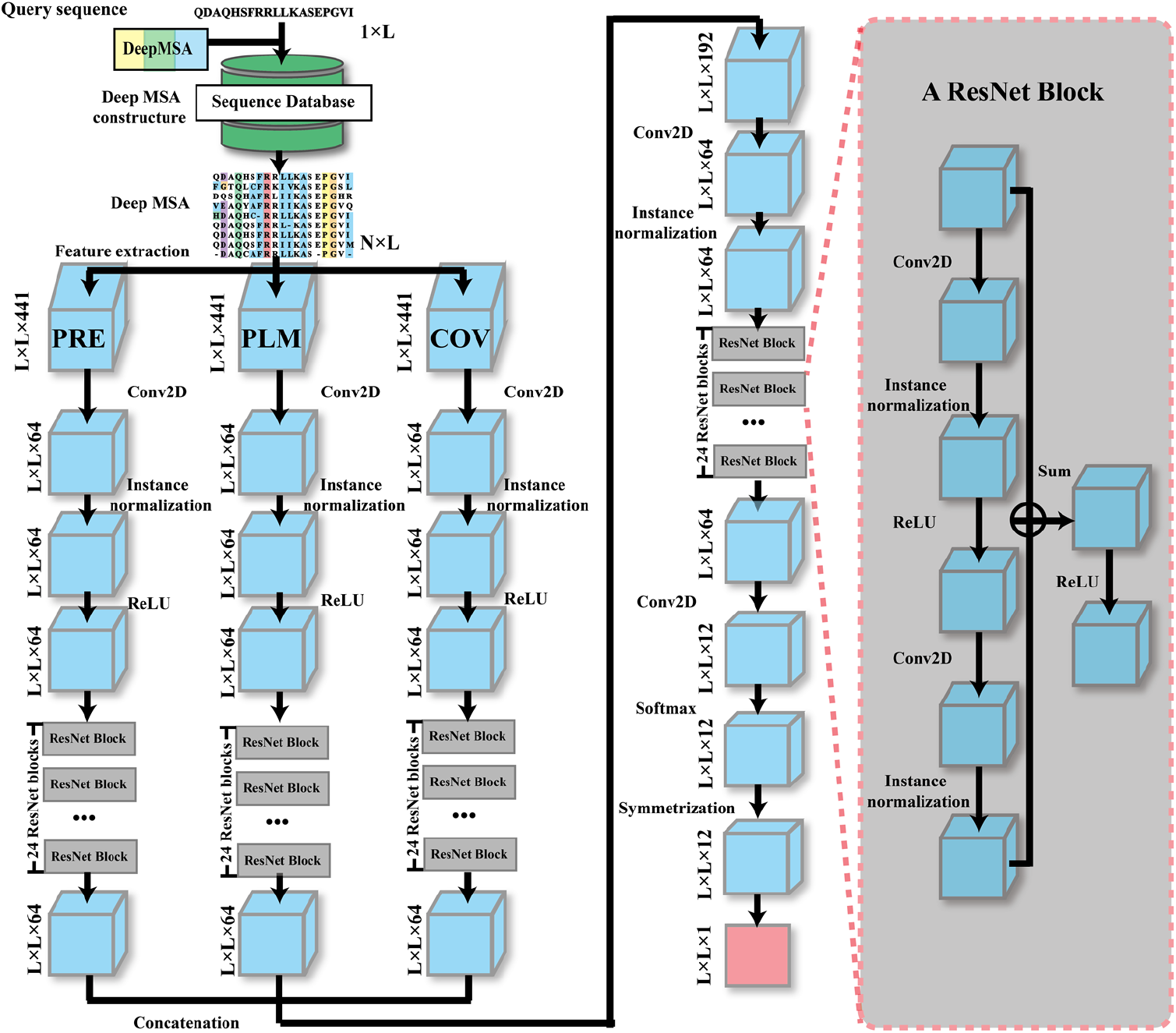
The architecture of TripletRes, which formulates the contact-map prediction as a pixel-level labeling problem, where a pixel in the image represents a pair of residue positions in the contact-map of the query protein. Starting from the MSA generated for the query sequence, three *L×L*×441 feature matrices (also called tensors) are computed for the three sets of coevolutionary features (PRE, PLM, COV). Here, *L* is the length of the query sequence while 441=21×21 is the combination of all 21 amino acid types (including the gap) for two positions in the MSA. Each tensor is input to a separate ResNet, where the first layer reduces the number of feature channels from 441 to 64, followed by instance normalization and 24 consecutive residual blocks to get an *L×L*×64 tensor. Details of a residual block are shown on the right-hand side inset. The three tensors from the three ResNets are concatenated into an *L×L*×192 tensor to feed into a final ResNet. In this ResNet, the first layer again reduces the feature channels from 192 to 64, followed by instance normalization, and 24 residual blocks to get an *L×L*×64 tensor, which is further reduced to *L×L*×12. Finally, a softmax layer is used to scale the values in the tensor between 0 and 1 and to make the sum of all values for each pixel (i.e. residue pair) equal to one. Since a protein contact/distance map is symmetric, TripletRes averages the corresponding softmax output of residue pair (*i,j*) and (*j,i*) to get the final *L×L*×12 distancemap prediction, where 12 stands of the number of distance bins. The contact-map is obtained by summing up the first 4 distance bin.

## Results and discussion

To examine the contact prediction pipelines, we collected two independent sets of test targets, including 50 nonredundant free-modeling (FM) domains from the CASP11 and CASP12 and 195 nonredundant targets assigned as *hard* by CAMEO(29). TripletRes was trained on 7,671 nonredundant domains collected from SCOPe-2.07(30). Detailed procedure to obtain the training and testing datasets are described in Text S1 in Supplementary Information, SI.

### Overall performance of TripletRes

Following the CASP criterion(4), two residues are defined as in contact if the Euclidian distance between their Cβ atoms (or Cα in case of Glycine) is below 8.0 Å. In this study, the accuracies, or mean precisions, of the top *L*/10, *L*/5, *L*/2, and *L* of medium- (12≤|*i − j*|≤23) and long-range (|*i − j*|≥24) contacts are evaluated, where *i* and *j* are sequential indexes for the pair of considered residues and *L* is the sequence length of the target. We focus on the performance on FM targets (or hard targets in CAMEO) and on long-range contacts for evaluation, since the metric is most relevant for assisting the prediction of the tertiary structure of non-homologous proteins(6, 31).

Table 1 summarizes the overall performance of long-range contact prediction on the two test datasets by TripletRes, in control with five state-of-the-art methods which are available for free-download and run with default setting (see Text S2 of SI for introduction of the control methods). The results show that TripletRes creates contact models with a higher accuracy than the control methods in all separation ranges for both test datasets. For example, on the 50 FM CASP targets, the average precision of the long-range top *L*/10, *L*/5, *L*/2, and *L* predicted contacts by TripletRes is 55.1%, 53.2%, 57.1%, and 58.4% higher, respectively, than the precision achieved by DeepContact, the most accurate third-party program in this comparison, which correspond to statistically significant *p*-values of 4.1e-08, 2.5e-07, 4.4e-10, and 1.1e-11 in the Student’s t-test. Notably, TripletRes only uses coevolutionary features, which is a subset of the diverse features employed by DeepContact. The better performance is probably due to the more effective integration of raw coevolutionary information in the TripletRes neural-network training.

**Table 1.**
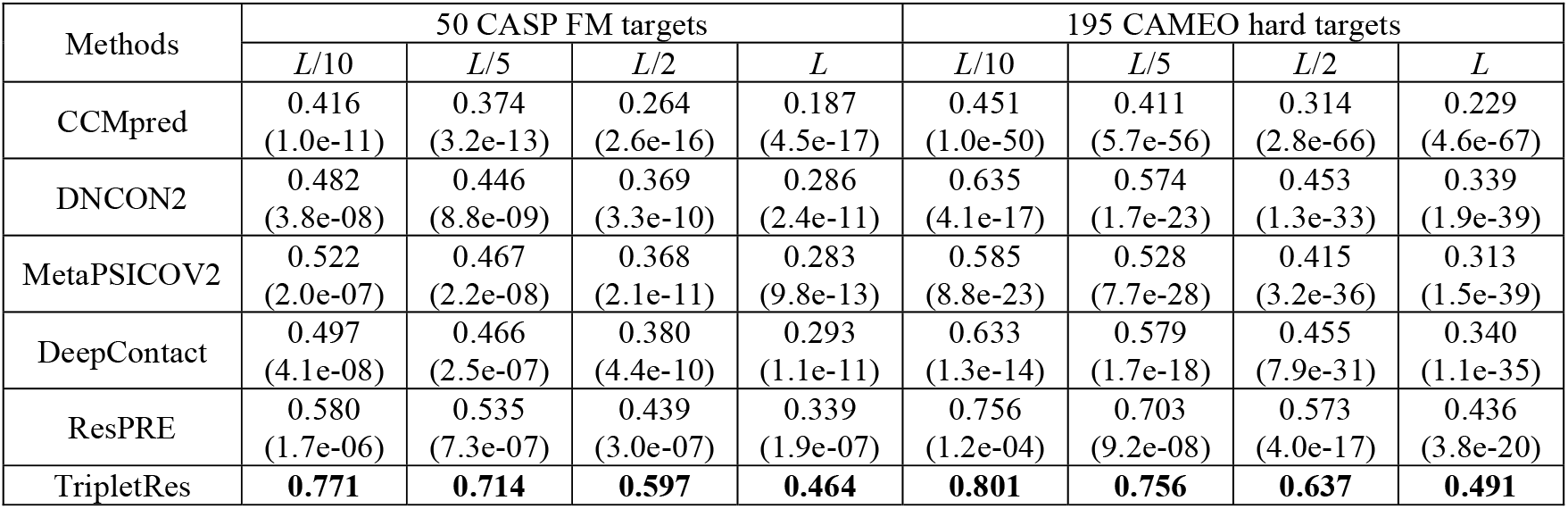
Summary of long-range contact precision by TripletRes and control methods on 50 CASP11&12 FM targets and 195 CAMEO hard targets, sorted in ascending order of top-*L* precision. *p*-values in parenthesis are from a Student’s t-test between TripletRes and each of the control methods, where bold fonts highlight the best performer in each category.

TripletRes also outperforms ResPRE, an in-house program previously trained on precision matrix(26), by a large margin. The long-range top-*L* precision of TripletRes is 36.9% higher than that of ResPRE with a *p*-value of 1.9e-07 on 50 FM targets. ResPRE achieved a significantly higher precision on CAMEO than the FM dataset, but its precision is still lower than that of TripletRes. For example, the mean precision of the top-*L* long-range contacts by TripletRes is 12.6% higher than that of ResPRE on the CAMEO targets. Given that both programs utilized the same precision matrix feature, the superiority of TripletRes is mainly attributed to the integrations of triplet coevolutionary features. In addition, as examined in detail below, the supervision of the distance predictions and the new deep MSA constructions also helped improve the accuracy of the TripletRes models.

### Feature extraction based on raw potentials outperforms that with post-processing

Feature extraction is essential for all machine-learning based modeling approaches. To quantitatively examine the effectiveness of the feature extraction strategy and the contribution of different feature types in TripletRes, we compare in Figs 2a-c the performance of two feature extraction strategies, based on three component features from covariance (COV), precision (PRE), and pseudolikelihood maximization (PLM) analyses (see Methods), respectively. The first feature extraction strategy, which was used by TripletRes, uses the raw coevolution potentials as input features, while the second strategy, which was commonly employed in many state-of-the-art predictors (21, 23, 24, 32), employs a specific post-processing procedure as described in Supplementary Eqs. S1 and S2 in SI Text S3. Since the traditional coevolutionary features can also be used to predict contacts directly without using supervised training, we list their performance as baselines (see dotted lines in Figs 2a-c). Here, a total of 767 sequences are randomly selected from 7,671 non-redundant SCOPe proteins as the validate set, while the remaining 6,904 sequences are used as the training set for feature extraction strategy selection in TripletRes. All experiments are performed by keeping other elements (e.g., MSA generation, neural network structure and its hyper-parameters) fixed.

**Fig 2.**
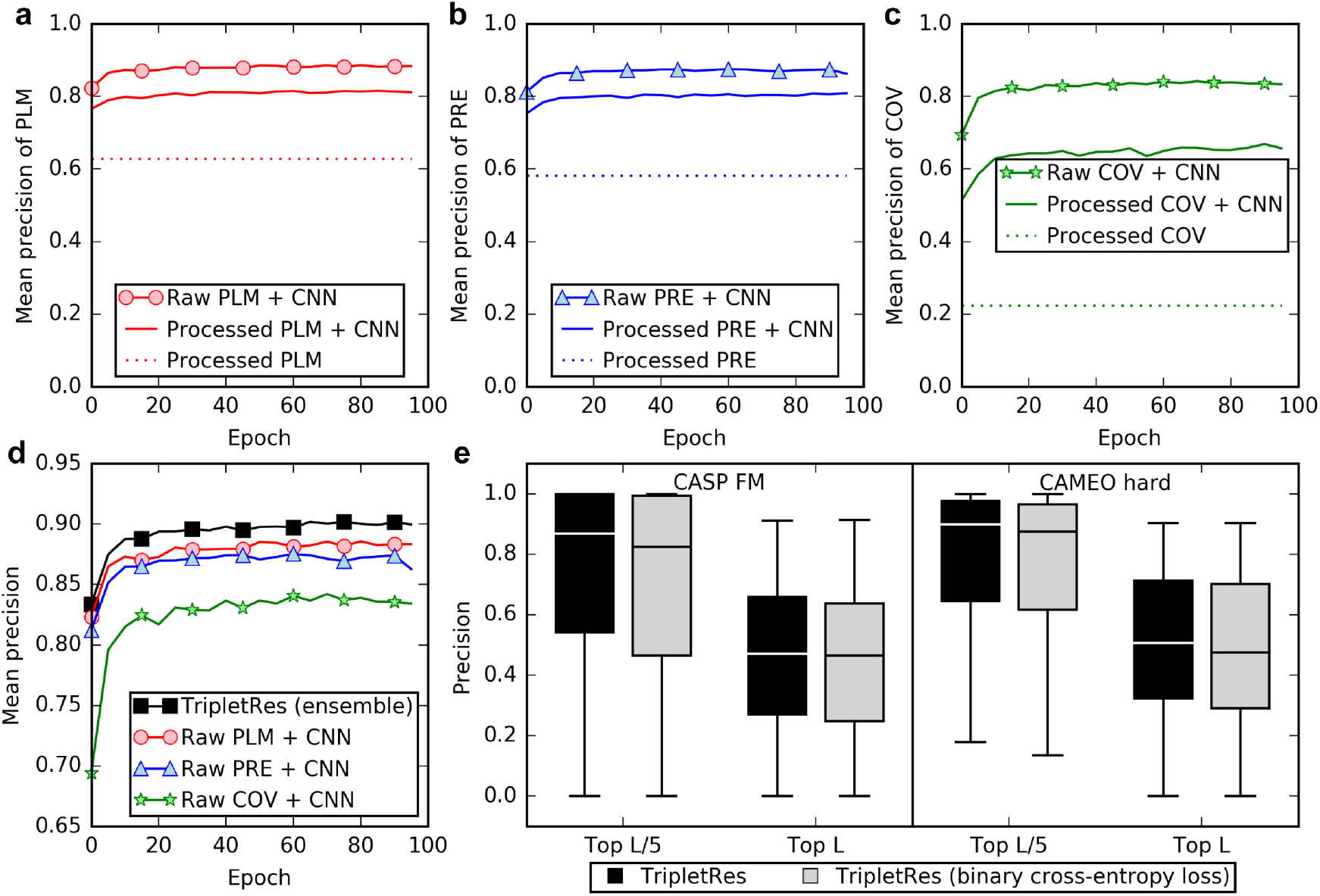
Comparisons of different strategies used to train TripletRes. **(a-c)** Comparisons of the average long-range top-*L*/5 precisions over training epochs using different feature extraction strategies but trained with the same deep neural-network structure on three different coevolutionary analysis methods: **(a)** DCA based on pseudolikelihood maximization (PLM), **(b)** DCA based on the precision matrix (PRE), **(c)** Covariance analysis (COV) for contact-map prediction, on the validation set. “Processed” means the coevolutionary features are post-processed by Eqs. S1 and S2 in SI Text S3. **(d)** Comparison of the average long-range top-*L*/5 precisions over training epochs of individual coevolutionary features and the TripletRes model that ensembles all three sets of features, on the validation set. **(e)** Comparison of long-range top-*L*/5 and top-*L* precisions with different loss functions on the CASP FM and CAMEO hard targets.

It can be observed from Figs. 3a-c that the new feature extraction strategy achieves a better contact prediction performance compared to the traditional feature extraction for all three considered matrix features. The highest mean precisions of the new feature extraction strategy on the long-range top-*L*/5 contact prediction are 84.2%, 87.5%, and 88.6%, respectively, for COV, PRE, and PLM features. If the post-processed features of Eqs. S1-2 are used, the mean precisions are reduced to 66.8%, 80.8%, and 81.6%, which represent a precision drop by 20.7%, 7.7%, and 7.9%, respectively, compared to the TripletRes feature extraction strategy. On the other hand, the mean precisions of both feature extraction strategies are consistently higher than the baseline through the training epochs, indicating the necessity of supervised training.

**Fig 3.**
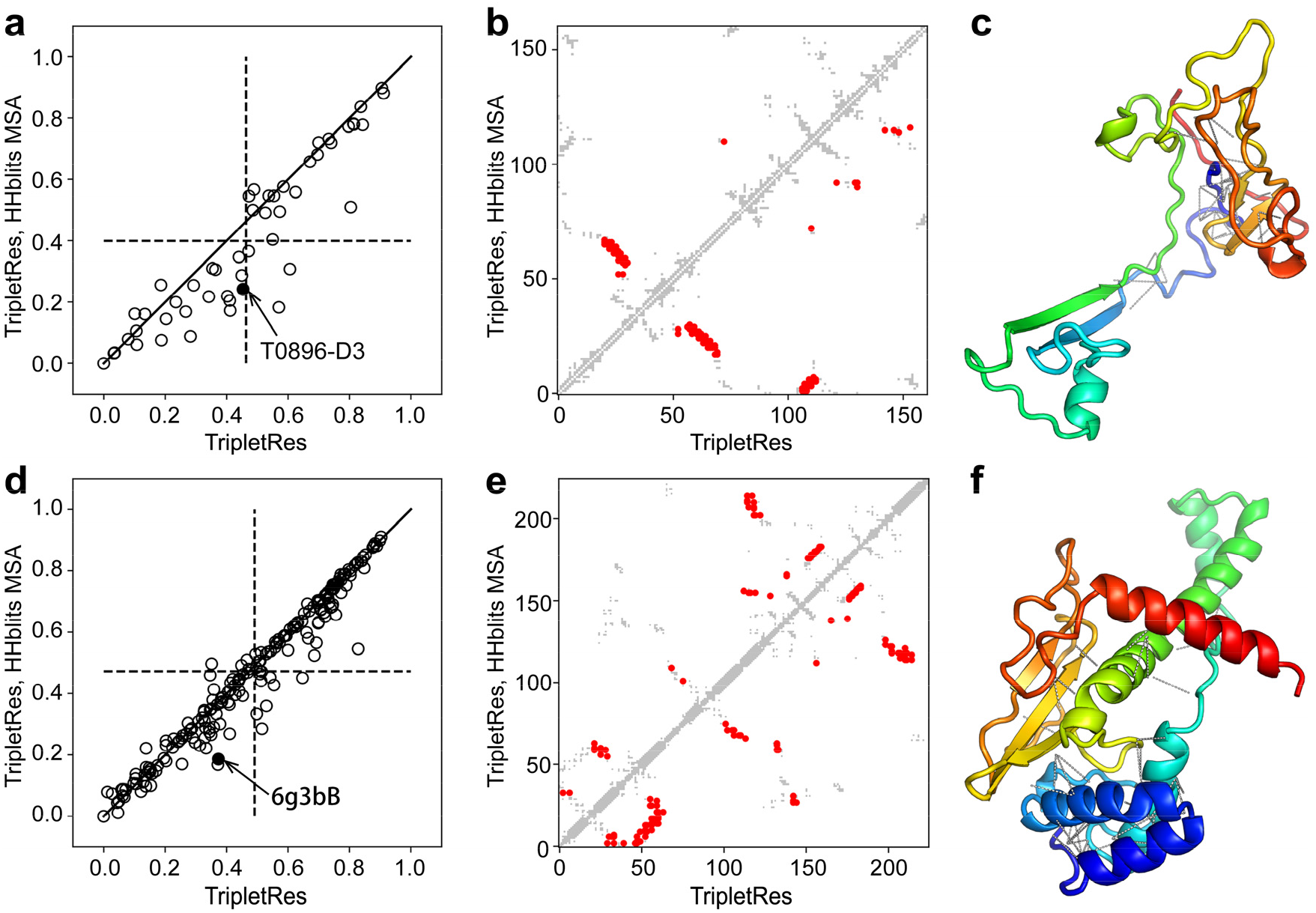
Long-range top-*L* precision of contact-maps predicted by TripletRes with deep MSAs versus that without deep MSAs. **(a)** overall results on 50 CASP FM targets; (**b,c**) illustrative example of contact-map and the native structure of the T0896-D3 domain in CASP12; (**c**) Overall results on 195 CAMEO hard targets; **(d,e)** illustrative example of the contact-map and the native structure of the PDBID 6g3bB in CAMEO. In (**a**) and (**d**), dashed lines mark the average precision of the top-L long-range contact prediction. In (**c**) and (**f**), dashed lines label the additional contacts predicted due to the employment of deep MSA.

One reason for the performance degradation by the post-processing approach is that the potential score for different types of residue-pairs have been treated equally and the sign of these potential scores is thus completely ignored in Eq. S1, when the post-processed coevolutionary features are fed to the supervised models. In contrast, the approach in TripletRes can keep detailed score information of different residue-pair types from the coevolutionary analyses for each residue pair, and thus allow for deep residual neural networks to automatically learn the inter-residue interactions not only on the spatial information but also on the residue pair-specific scores of different residue-pair types, while the traditional supervised machine learning models can only learn the spatial information of each residue pair during the training.

### Ensembling different component features improves contact-map prediction

Compared to ResPRE(26), a major new development in TripletRes is on the integration of multiple coevolutionary feature extractions. To examine the efficiency of ensembled feature collection on the contact predictions, Fig 2d presents the average long-range top *L*/5 precisions of the predictors trained by three individual component features and their ensemble. All models shown in Fig 2d become stable after 40 rounds of training, and obtain a precision of 88.4%, 86.3%, 83.4%, and 90.0% when using PLM, PRE, COV and an ensemble of all three features, respectively, after 100 epochs of training. In general, the COV-based model has the lowest precision among the three individual feature models, probably due to the translational noise in the covariance matrix(26). The performance of the two DCA-based features by PRE and PLM are comparable and both consistently outperform COV by a large margin. TripletRes ensembles three features that can obtain more comprehensive coevolutionary information from the deep MSAs. As a result, the ensemble model has a higher precision than all models from the component features, demonstrating the effectiveness of multiple feature integration.

### Loss function with continuous distances outperforms that with binary contacts

The correct loss function selection plays an important role in the training of neural networks because it determines the performance metric of the model during training. The most commonly used loss function for contact-map predictions is the binary cross-entropy loss function, which encodes each residue pair with 2 states (contact and not in contact). Typically, with a single distance threshold of 8 Å, such a loss function does not encode detailed distance information, e.g., residue pairs separated by 9 Å will be treated the same as those by 22 Å. In contrast, the loss function in TripletRes (Eq. 6 in Methods) considers a discrete representation of the supervised information for each residue pair and thus could keep more distance information for training.

Fig 2e compares the long-range top-*L*/5 and top-*L* precisions between TripletRes programs trained with Eq. 6 or a binary cross-entropy loss (see Eq. S3 in SI Text 4) on CASP FM and CAMEO hard targets, respectively. It can be observed that incorporating continuous distance information in training can lead to improvements in contact-map prediction, even though the contact-maps are not directly optimized. For example, the distance information in the loss function can improve the top *L*/5 precision from 68.7% to 71.4%, and 74.1% to 75.6%, for the CASP set and CAMEO set, respectively, which correspond to a *p*-value of 2.1e-02 and 7.9e-03 in Student’s t-test. Interestingly, when more top-ranked contacts are considered (i.e., top *L*), the *p*-values becomes more significant and decrease to 1.6e-04 and 2.0e-07 on the two datasets, respectively, which means the distance information may have a stronger effect on improving the precision when more contacts are evaluated. Protein structure prediction methods can thus benefit more from TripletRes, which was trained with the discrete distance loss function because more predicted contacts can be reliably considered as restraints for protein folding.

### Deep MSA search help create more comprehensive coevolutionary information

TripletRes utilizes MSAs as the only input and the quality of the latter is thus essential to the final contact prediction models. It is worth noting that the TripletRes model is trained on features extracted from MSAs generated by HHblits, but a deeper MSA generated by multiple databases (Methods) has been used for test proteins. We expect the strategy could reduce over-fitting between the training and test proteins.

To examine the impact of different MSA collections on the contact models, Fig 3 shows a comparison of TripletRes models with and without deep MSAs on the test proteins from CASP FM targets (Fig 3a) and CAMEO hard targets (Fig 3d). Here, dashed lines mark the mean precision value of the long-range top-*L* prediction by each dataset. For the CASP FM targets, the usage of deep MSAs during testing significantly improves the mean precision of TripletRes from 40.0% to 46.4% with a *p*-value 1.9e-05 in Student’s t-test, where 35 out of 50 FM targets (70%) achieve a higher precision with deep MSAs while only 8 targets (16%) do so when the HHblits MSAs are used. The same trend can be observed in the CAMEO targets, where the *p*-value of improvement in long-range top-*L* precision is 1.7e-06. This difference is mainly due to the higher number of homologous sequences collected in deep MSA search protocol, which allows the extraction of more reliable coevolutionary information. For example, the average number of effective sequences of MSAs, or Neff calculated by Eq. 1 in Methods, generated by deep MSA is 85.4, which is 34.3% higher than that obtained by HHblits on CASP FM targets (63.6).

In Figs 3b-c and Figs 3e-f, we select two illustrative cases from the CASP and CAMEO datasets respectively. The example in Fig. 4b is from the third domain of CASP12 target T0896 with experimental structure presented in Fig. 4c, where HHblits collects a relatively shallow MSA with a Neff=0.94, which resulted in only 39 true positives in the 162 long-range top-*L* contact predictions. The deep MSA search increased the Neff value to 3.78, where the number of true contacts with the deeper MSA increases to 73, which is 87.2% higher than that with the HHblits MSA. In Figs 3e-f, the structure comes from the Type II site-specific deoxyribonuclease (PDBID: 6g3bB) with 225 residues, where HHblits creates an MSA with Neff=2.0 and results in 42 true positives out of top-*L* long-range predictions; while 42 more contacts are detected by TripletRes through the deep MSA that has a Neff of 14.0. These examples highlight again the importance of using deep MSA pipeline for coevolutionary feature collection and the impacts on final contact-map prediction. The new contacts correctly predicted after performing deep MSA searching strategy are marked as dashed lines in Fig 3c and 3f; these contacts provide additional spatial restraints and have shown critical in creating correct global fold for the domain structures(33).

**Fig 4.**
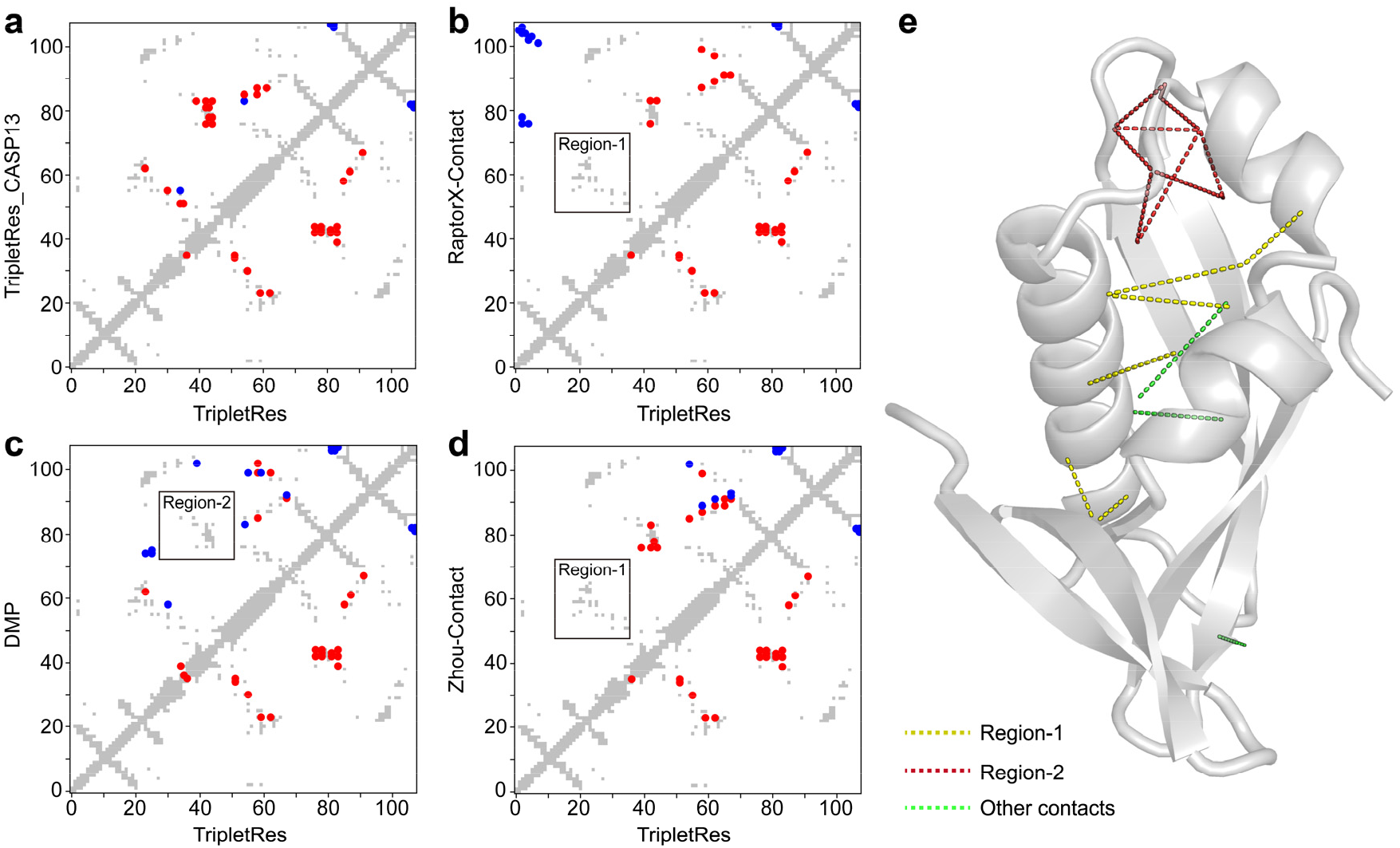
An illustrative example of a CASP13 domain T0957s1-D1 showing a comparison of top-*L*/5 long-range contact prediction by TripletRes and the control methods. In each map, the true contacts are marked in grey, true positives in red, and false positives in blue. **(a-d)** The comparison between TripletRes_CASP13, RaptorX-Contact, DMP, and ZHOU-Contact (in upper-left triangle) against TripletRes (in lower-right triangle). **(e)** Experimental structure of T0957s1-D1, with the long-range true positive prediction by TripletRes in Region 1, Region 2 and others marked in yellow, magenta and green dashed lines, respectively.

### Performance of TripletRes for blind prediction in CASP13

An early version of TripletRes, denoted as TripletRes_CASP13, participated in the 13th CASP experiment for inter-residue contact prediction(6, 34). It was ranked among the top two methods based on the mean precision score (http://www.predictioncenter.org/casp13/zscores_rrc.cgi), with another top method RaptorX-Contact which also ranked as the top method in previous CASPs. In Table 2, we list a summary of the average results by TripletRes and TripletRes_CASP13, along with three other top CASP13 predictors from RaptorX-Contact, DMP, and ZHOU-Contact. For the long-range top-*L*/5 contacts on the 31 FM targets, TripletRes_CASP13 achieved a mean precision of 64.6%, while the mean precision of RaptorX-Contact, DMP, and ZHOU-Contact are 69.4%, 60.2%, and 58.3%, respectively. TripletRes, however, achieves the highest precision of 71.6% for long-range top *L*/5 contacts. Here, the major difference between TripletRes and TripletRes_CASP13 is that TripletRes utilizes a new loss function (Eqs. 6-7 in Methods) to integrate distance profiles for contact-maps, while TripletRes_CASP13 used a binary cross-entropy loss function (Eq. S3 in Text S4 of SI). These data demonstrate the validity of the distance-supervised training strategy.

**Table 2.** Performance comparisons on CASP13 FM targets between TripletRes and RaptorX-Contact, DMP, and ZHOU-Contact servers, sorted in ascending order of top *L* long-range contact precision.

| Method | Medium range |  |  |  | Long-range |  |  |  |
| --- | --- | --- | --- | --- | --- | --- | --- | --- |
| | $L/10$ | $L/5$ | $L/2$ | $L$ | $L/10$ | $L/5$ | $L/2$ | $L$ |
| ZHOU-Contact | 0.727 | 0.623 | 0.453 | 0.319 | 0.641 | 0.583 | 0.474 | 0.367 |
| DMP | 0.772 | 0.682 | 0.505 | 0.344 | 0.645 | 0.602 | 0.470 | 0.361 |
| TripletRes CASP13* | 0.842 | 0.746 | 0.543 | 0.360 | 0.695 | 0.646 | 0.534 | 0.409 |
| RaptorX-Contact | 0.805 | 0.702 | 0.527 | 0.364 | 0.762 | 0.694 | 0.567 | 0.438 |
| TripletRes* | **0.865** | **0.770** | **0.562** | **0.367** | **0.775** | **0.716** | **0.573** | **0.440** |

\* “TripletRes” is the current version of TripletRes trained using distance-based loss function (Eq. 6). “TripletRes\_CASP13” is the early version of TripletRes used in CASP13, trained using binary cross-entropy loss function (Eq. S3 in Text S4 of SI).

In Fig 4, we present an example from the first domain of T0957s1 of CASP13 which is a contact-dependent growth inhibition toxin-immunity protein (PDB ID:6cp8) with an α+β fold and 108 residues. TripletRes collected a deep MSA with Neff=6.7, significantly higher than the Neff value (1.3) by HHblits. This resulted in a mean precision of 86.4% for the top-*L*/5 long-range contact predictions, compared to 40.9% by RaptorX-contact, 36.4% by DMP, and 54.5% by ZHOU-Contact, respectively. TripletRes also performed better than the CASP13 version in precision (77.3%), benefited from the distance information during the training. As shown in Figs 4b and 4d, RaptorX-Contact and ZHOU-Contact failed to hit any long-range contacts in Region 1 which is a critical loop-loop contact region. DMP, on the other hand, was not able to cover contacts in Regions 2 that are important to pack the core structure of the two helices with the center beta-sheet (Fig 4c). TripletRes can cover both Regions marked in yellow and magenta in Fig 4e, respectively. Among the top-*L*/5 correctly predicted long-range contacts, 94.7% of them have the distance profile with a probability peak at <8Å and nearly 74% of the residue pairs have the accumulated probability >80% in the region below 15Å, indicating a high confidence of contact prediction on the residue pairs based on the distance profile.

## Conclusion

Protein contact-map prediction has been critical to assist protein folding in the form of spatial constraints. This work presented a new deep learning method for high-accuracy contact prediction by learning from raw coevolutionary features extracted with deep multiple sequence alignments. The method was tested on FM domains in CASP11-13 and hard targets from CAMEO experiments, which demonstrated a significant advantage in accurate contact-map predictions compared to other state-of-the-art approaches.

Several factors were found to contribute to the success of the TripletRes pipeline. First, coupling deep residual convolutional networks directly with raw coevolutionary matrices can result in better performance than feeding neural networks with the post-processed features. Second, a triplet of coevolutionary features, from covariance matrix, inverse covariance matrix and the inverse Potts model approximated by pseudolikelihood maximization, are ensembled in TripletRes by a set of four neural networks constructed with residual blocks. This feature ensemble strategy was found to enable more accurate prediction than using the three sets of features individually. Third, including more discrete distance information into the network training was proven to be beneficial to the contactmap prediction compared to binary contact training, although the contact-map models are binary on their own. This is largely because the distance-based loss function enables the learning of detailed spatial features specified by the sequence profiles. Finally, a hierarchical sequence searching protocol was proposed to obtain deeper MSAs, which impact the performance of the final model prediction.

It is worth noting that the major goal of contact-map prediction is for assisting *ab initio* 3D structure construction, where a significant amount of efforts has been made along this line in the past decades(8, 31, 35-37). Although recent progress of the field has shown an advantage of distance predictions(32, 38), contact-map can provide reliable information of short-distance residue-residue interactions that is critical to specify the global topology of the protein fold. In fact, our results showed that most of the accurately predicted distances in TripletRes are still on the residues pairs with a short distance below 9-10 Å, which is part of the reason that has motivated our idea of distance-supervised learning in TripletRes. In addition, the development of feature extraction for protein contact-map prediction has direct contributions to the prediction of other forms of long-range residue-residue interactions. Therefore, with the development of new approaches and consistent improvement of the model accuracy, the advanced sequence-based contact-map predictions will continue to be an important driving force for template-free structure prediction of the field.

## Methods and Materials

TripletRes is a deep-learning based contact-map prediction method consisting of three consecutive steps (Fig 1). It first creates a deep MSA and extracts three coevolutionary matrix features. Next, the feature sets are fed into three sets of deep ResNets and trained in an end-to-end fashion. Finally, a symmetric matrix distance histogram probability is created and binarized into the contact-map prediction.

### MSA generation

To help offset the overfitting effects, TripletRes creates MSAs using different strategies for training and testing protein sequences. For training proteins, MSAs are created by HHblits with an E-value threshold of 0.001 and a minimum sequence coverage of 40% to search through the latest Uniclust30(39) database with 3 iterations.

For test proteins, the initial MSA is created also by HHblits but followed-up with multiple iterations. If the Neff value of the initial MSA is lower than a given threshold (=128 that was decided by trial and error), a second step will be performed using jackhmmer(40) through UniRef90(41). Here, Neff measures the number of effective sequences in the MSA and is defined as:

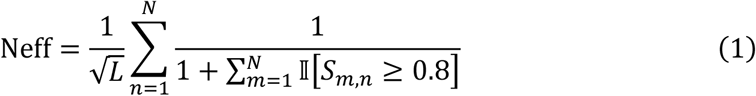

where *N* is the total number of sequences in the MSA, 𝕀[*S_m,n_* ≥ 0.8] = 1 if the sequence identity *S_m,n_* between sequences *m* and *n* is over 0.8; or = 0 otherwise. To assist the MSA concatenation, the jackhmmer hits are converted into an HHblits format sequence database, against which a second HHblits search was performed. In case that Neff is still below 128, a third iteration is performed by hmmsearch(40) through the MetaClust(42), where the final MSA is pooled from all iterations (see Fig S1 in SI for the whole MSA construction pipeline).

### Coevolutionary feature extraction

Three sets of coevolutionary features are extracted from the deep MSAs. First, the covariance (COV) feature measures the marginal dependency between different sequential positions and is calculated by

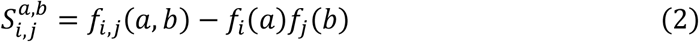

where *f_i_*(*a*) is the frequency of a residue *a* at position *i* of the MSA, *f_i,j_*(*a, b*) is the cooccurrence of two residue types *a* and *b* at positions *i* and *j*.

The second precision matrix (PRE) feature, Θ, which represents an inverse of the covariance matrix *S*, can be estimated by minimizing the regularized negative log-likelihood function(34)

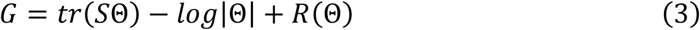

where the first two terms are the negative log-likelihood of Θ assuming that the data follows a multivariate Gaussian distribution; *tr*(*S*Θ) is the trace of matrix *S*Θ; *log*|Θ| is the log determinant of Θ; and 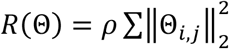 is the regularization function of Θ to avoid over-fitting, with *ρ = e*^−6^ being a positive regularization hyper-parameter.

The last feature is a coupling parameter matrix of inverse Potts model approximated by PLM. Instead of assuming the data follows a multivariate Gaussian distribution, PLM approximates the probability of a sequence for the Potts model with

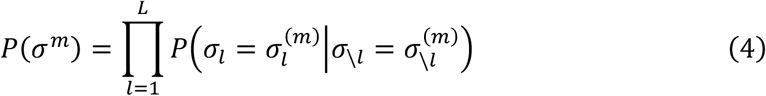

Here, *P*(*σ^m^*) is the probability model for the *m*-th sequence in the MSA and 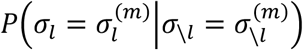 is the marginal probability of *l*-th position in the sequence by

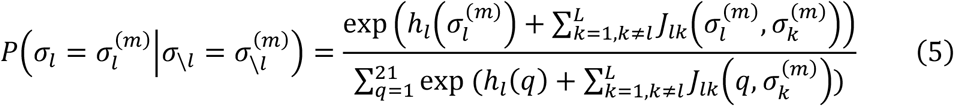

where *h* and *J* are single site and coupling parameters, respectively. In TripletRes, the raw coupling parameter matrix *J* is used as the PLM feature.

Thus, each feature is represented by a 21 ∗ *L* by 21 ∗ *L* matrix for a protein sequence with *L* amino acids. The entries of the 21 by 21 sub-matrix of a corresponding amino acid pair are the descriptors, which are fed into a convolutional transformer as conducted by a fully convolutional neural network with residual architecture (Fig 1).

### Deep neural-network modeling

TripletRes implements residual neural networks (ResNets)(28) as the deep learning model. Compared to traditional convolutional networks, ResNets adds feedforward neural networks to an identity map of input, which helps enable the efficient training of extremely deep neural networks such as the one used in TripletRes. As illustrated in Fig 1, the neural network structure of TripletRes has four sets of residual blocks, where three of them are connected to the input layer for feature extraction. Each of the three ResNets has 24 basic blocks and can learn layered features based on the specific input. After transforming each input feature into a feature map of 64 channels, we concatenate the transformed features along the feature channel and employ another deep ResNet containing 24 residue blocks to learn the fused information from the three features.

The activation function of the last layer is a softmax function which outputs the probability of each residue pair belonging to specific distance bins. Here, the residueresidue distance is split into 10 intervals spanning 5-15Å with an additional two bins representing distance less than 5Å and more than 15Å, respectively. The whole set of deep ResNets are trained by the supervision of the maximum likelihood of the prediction, where the loss function is defined as the sum of the negative log-likelihood over all the residue pairs of the training proteins:

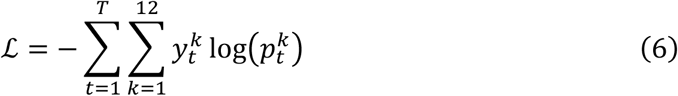

Here, *T* is the total number of residue pairs in the training set. 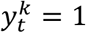 if the distance of *t*-th residue pair of native structures falls into *k*-th distance interval; otherwise *y^k^* = 0. 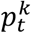 is the predicted probability that the distance of the *t*-th residue pair falls into the *k*-th distance interval. The probability of the *t*-th residue pair forming a contact *P_t_* is the sum of the first 4 distance bins:

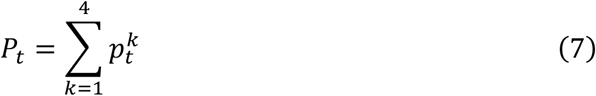

The training process uses dropout to avoid over-fitting, where the dropout rate is set to 0.2. We use Adam(43), an adaptive stochastic gradient descent algorithm, to optimize the loss function. TripletRes implements deep ResNets using Pytorch(44) and was trained using the Extreme Science and Engineering Discovery Environment (XSEDE)(45).

## Acknowledgements

This work used the Extreme Scienceand Engineering Discovery Environment (XSEDE), which is supported by National Science Foundation (ACI-1548562). The work was done when Yang Li visited at University of Michigan.

## Author contributions

Y.Z. conceived the project; Y.L. developed the method and performed the experiment; C.Z. developed deep MSA pipeline; Y.L. and C.Z. developed the webserver; E.W.B., W.Z., X. Z analyzed the data and participated in discussions; D.Y. and Y.Z. supervised the project; Y. L. and Y.Z. wrote the manuscript; all authors proofread and approved the manuscript.

## Supporting Information

### Supporting Figures

- Figure S1. The DeepMSA pipeline for generating deep multiple sequence alignments for TripletRes.

### Supporting Texts

- Text S1. Detailed procedure to collect training and test datasets.
- Text S2. A brief introduction of control methods and other top participants in CASP13.
- Text S3. Traditional feature extraction strategy with post-processing.
- Text S4. Binary cross entropy loss function for training TripletRes in CASP13.

