## Supplementary Material for "Deducing high-accuracy protein contact-maps from a triplet of coevolutionary matrices through deep residual convolutional networks"

### Supporting Information

#### Supporting Figures

- Figure S1. The DeepMSA pipeline for generating deep multiple sequence alignments for TripletRes.

#### Supporting Texts

- Text S1. Detailed procedure to collect training and test datasets.
- Text S2. A brief introduction of control methods and other top participants in CASP13.
- Text S3. Traditional feature extraction strategy with post-processing.
- Text S4. Binary cross entropy loss function for training TripletRes in CASP13.

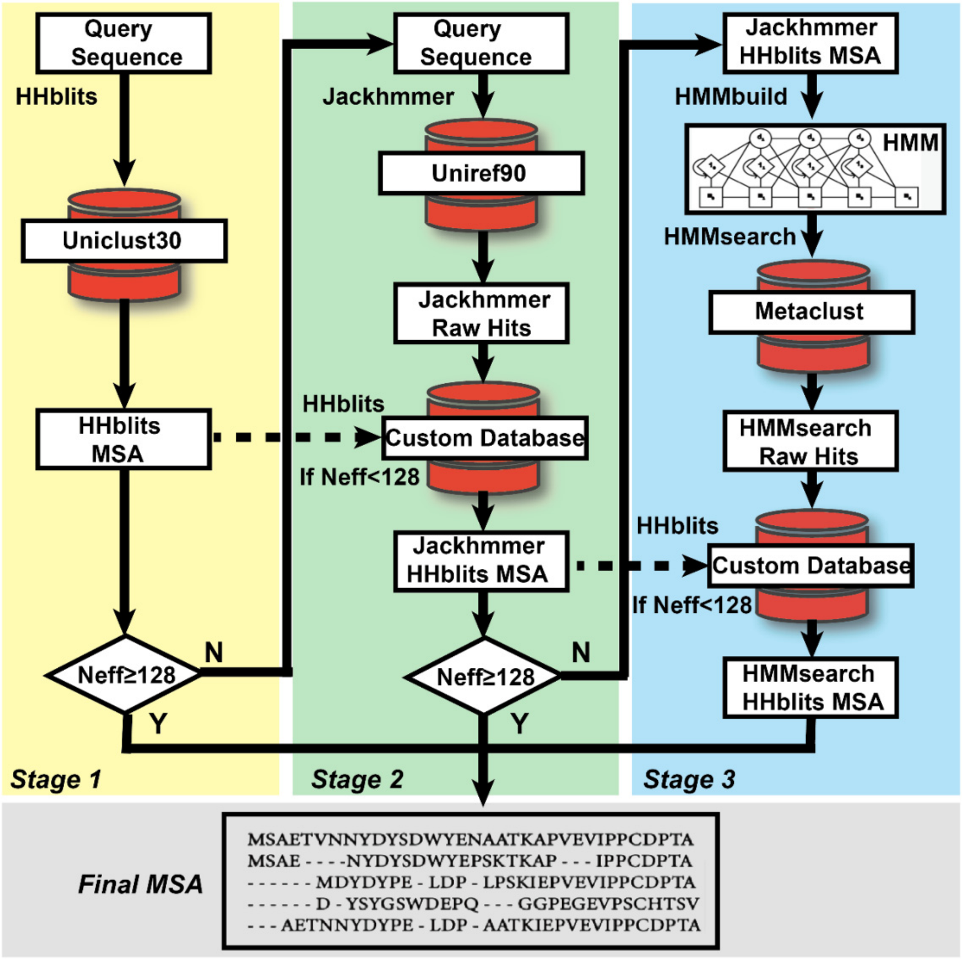


**Fig S1.** The DeepMSA pipeline for generating deep multiple sequence alignments for TripletRes. DeepMSA consists of three stages. The query sequence is first searched by HHblits against the Uniclust30 database to generate Stage 1 MSA (yellow background). In Stage 2 (green background), Jackhmmer searches the query sequence through the UniRef90 database to find sequence homologs, which are built into a custom database in HHblits format. HHblits is then used to search Stage 1 MSA through this custom database to get Stage 2 MSA. In Stage 3 (cyan background), the Stage 2 MSA is converted into a hidden Markov model (HMM) by HMMbuild and used by HMMsearch to search the Metaclust metagenome protein sequence database. The identified sequence homologs are reformatted to another HHblits format custom database. The MSA from Stage 2 is then used to search against this custom database to get the final stage alignment. In this incremental MSA construction process, if the MSA from either Stage 1 or Stage 2 reaches Neff>=128, this MSA will be output as final MSA without subsequent stages.

**Text S1. Detailed procedure to collect training and test datasets**

The 50 nonredundant FM domains from the CASP11 and CASP12 experiments are downloaded from http://predictioncenter.org/download_area/. The 195 nonredundant targets defined as *hard* by CAMEO are collected from https://www.cameo3d.org/sp/targets/1-year/?to_date=2019-07-20 whose Submission Date is from 2018-07-28 to 2019-07-20. Both of the two test datasets have a pair-wise sequence identity < 30% within each of the two datasets. All sequences share >30% to any proteins used in training the TripletRes models are removed. For a fair comparison, those targets in test sets were also removed if they cannot be finished in 72 hours by any of the control methods.

TripletRes was trained on a subset of SCOPe 2.07 domain sequences collected as per the following criteria: (1) Sequence length should be in the range of 30-400 residues; (2) Resolution of the corresponding structure should be better than 2.0 Å; (3) Maximum pairwise sequence identity is also set to 30%. There were 7,671 domains collected for training. The whole training set was split into 10 subsets, and we randomly selected one as the validation set and left the remaining subsets as the training set for hyper-parameter tuning. After the hyper-parameter tuning, the final model is the average of 10 models and each model was trained by considering each subset as the validation set and the remaining subsets as the training set.

**Text S2. A brief introduction of control methods and other top participants in CASP13**

CCMpred[1] is a representative DCA method, and the several of other control methods, DNCON2[2], MetaPSICOV2[3], DeepContact[4], and ResPRE[5], are based on supervised machine learning models using outputs of CCMpred or other DCA methods as input feature. Here, DNCON2, MetaPSICOV2 and DeepContact are the top-ranking predictors in CASP12. ResPRE was our previous work which was built on raw precision matrix feature and shown to be comparable with many state-of-the-art methods despite the use of a single precision feature matrix. It should be noted that CCMpred does not have a built-in program for MSA generation. For a fair comparison, we tested it with the same MSAs as those used in the test phase of TripletRes. The control methods were downloaded and implemented in our local computers with default parameters.

In CASP13, DMP, also known as DeepMetaPSICOV[6], combines the input features of MetaPSICOV2 and a covariance feature[7] with residual convolutional neural networks (RCNNs). Meanwhile, both ZHOU-Contact, i.e. SPOT-Contact[8], and RaptorX-Contact[9] combine traditional one-dimensional features (secondary structure, solvent accessibility, and sequence profile, etc.) and pairwise coevolution features (CCMpred final output) by RCNNs or recurrent neural networks. The prediction results of other participants in CASP13 were obtained from CASP13 data archive.

**Text S3. Traditional feature extraction strategy with post-processing**

As a baseline for comparison, the traditional feature extraction method, which involves a post-processing procedure over a raw coevolutionary feature matrix, is tested. There are usually two steps in the post-processing procedure. The coevolutionary feature matrix is first transformed into an *L* by *L* contact score matrix *C* by

$$C_{ij} = \sqrt{\sum_{a,b} \left\| F_{ij}^{ab} \right\|_2^2} \tag{S1}$$

where each entry represents the potential of forming a contact. Here, *a* and *b* represent two types of amino acids, and *F* represents the obtained coevolutionary feature matrix. The contact score matrix *C* will be further normalized by an average product correction (APC) step:

$$C_{ij}^{APC} = C_{ij} - \frac{C_i C_j}{C} \tag{S2}$$

where $C_i = \frac{1}{L}\sum_{j \neq i}^{L} C_{ij}$, and $\mathrm{C} = \frac{1}{L^2 - L}\sum_{i,j,i<j}^{L} C_{ij}$. $C^{APC}$ is the predicted contact-map based on coevolution analysis with the post-processing procedure. $C^{APC}$ can be considered as the input feature of a supervised machine learning model. In this work, we use the same neural network structure with 22 residual blocks as the supervised learning model for the comparison of the two extraction strategies.

**Text S4. Binary cross entropy loss function for training TripletRes in CASP13.**

The loss function is defined as the sum of cross entropy over all the residue pairs of the training proteins:

$$\mathcal{L}_{bin} = -\sum_{t=1}^{T} y_t \log(p_t) + (1 - y_t) \log(1 - p_t) \quad \text{(S3)}$$

Here, *T* is the total number of residue pairs in the training set. $y_t = 1$ if the distance of *t*-th residue pair of native structure is below 8Å; otherwise $y_t = 0$. $p_t$ is the predicted probability that the *t*-th residue pair forms a contact.
